# Simultaneous auditory agnosia: Systematic description of a new type of auditory segregation deficit following a right hemisphere lesion

**DOI:** 10.1101/2020.02.27.968107

**Authors:** Emma Holmes, Nattawan Utoomprurkporn, Chandrashekar Hoskote, Jason D. Warren, Doris-Eva Bamiou, Timothy D. Griffiths

**Author notes:** Corresponding author: Emma Holmes; Mailing address: Wellcome Centre for Human Neuroimaging, UCL Queen Square Institute of Neurology, University College London, 12 Queen Square, London WC1N 3BG, U.K.

## Abstract

We investigated auditory processing in a young patient who experienced a single embolus causing an infarct in the right middle cerebral artery territory. This lead to damage to auditory cortex including planum temporale that spared medial Heschl’s gyrus, and included damage to the posterior insula and inferior parietal lobule. She first reported difficulty hearing all sounds, which fully recovered within days, but she subsequently reported chronic difficulties with segregating speech from noise and segregating elements of music. Clinical tests showed no evidence for abnormal cochlear function. Follow-up tests confirmed difficulties with auditory segregation in her left ear that spanned multiple domains, including speech-in-noise and music streaming. Testing with a stochastic figure-ground task—a way of estimating generic acoustic foreground and background segregation—demonstrated that this was also abnormal. This is the first demonstration of an acquired deficit in the segregation of complex acoustic patterns due to cortical damage, which we argue is a causal explanation for the symptomatic deficits in the segregation of speech and music. These symptoms are analogous to the visual symptom of simultaneous agnosia. Consistent with functional imaging studies on normal listeners, the work implicates non-primary auditory cortex. Further, the work demonstrates a (partial) lateralisation of the necessary anatomical substrate for segregation that has not been previously highlighted.

**Highlights:**

- Rare case of auditory agnosia in a young patient with a right-hemisphere infarct
- Damage affecting non-primary auditory cortex, but sparing primary auditory cortex
- Generalised auditory segregation deficit, revealed by auditory figure-ground task
- This explains segregation deficits for speech-in-noise and music streaming
- The deficit affects stimuli presented on the left

## 1. Introduction

In our everyday lives, we are often in environments that contain multiple competing sounds—from the sound of someone’s voice in a noisy café, to a violin melody that emerges from a large orchestra. The auditory system faces the challenge of parsing these sounds, so that we can focus on the voice of a particular person or a particular melody that we wish to hear out. Yet, we do not fully understand which brain regions are required to carry out these processes. Here, we report a rare case of a young patient who experienced a right hemisphere infarct and subsequently reported difficulty listening in environments containing multiple sounds, such as understanding speech in noisy places and picking out melodies in music.

Understanding speech when competing sounds are present (“speech-in-noise perception”) is particularly difficult for people with sensorineural hearing loss (Dubno, Dirks, & Morgan, 1984; Gatehouse & Noble, 2004; Helfer & Freyman, 2008). Yet, difficulties with speech-in-noise perception cannot be fully accounted for by the pure-tone audiogram, which is the most common clinical measure of peripheral hearing ability: Even people who perform normally on clinical tests of peripheral auditory function frequently visit the clinic reporting difficulties understanding speech in noisy places (Cooper & Gates, 1991; Hind et al., 2011; G. Kumar, Amen, & Roy, 2007). Sub-clinical variability in pure-tone thresholds has been estimated to account for approximately 15% of the variance in speech-in-noise performance among people (Holmes & Griffiths, 2019), meaning that the remainder of the variance must originate from other processes.

Central processes are likely to affect the ability to parse target speech from simultaneously occurring background sounds. Holmes and Griffiths (2019) found that fundamental auditory grouping processes—assessed by an abstract figure-ground task—helped to explain variability in speech-in-noise perception after accounting for the audiogram. Using functional magnetic resonance imaging (fMRI), they showed that fundamental grouping processes relevant to speech-in-noise perception depend on processes in auditory cortex (Holmes, Zeidman, Friston, & Griffiths, 2019). This is broadly consistent with studies showing activation of auditory cortex during speech-in-noise perception (Davis, Ford, Kherif, & Johnsrude, 2011; Eckert, Teubner-Rhodes, & Vaden, 2016; Kamourieh et al., 2015; Wong & Parrish, 2008), and also with studies of figure-ground segregation that show activity in planum temporale (Teki et al., 2016; Teki, Chait, Kumar, von Kriegstein, & Griffiths, 2011).

Although many studies have examined the neural basis of music perception and disorders of this (for reviews, see Clark, Goldren, & Warren, 2015; Griffiths, Rees, & Green, 1999; Peretz & Zatorre, 2005; Stewart, Von Kriegstein, Warren, & Griffiths, 2006), fewer have focussed on the ability to separate (“hear out”) a target melody in a musical piece containing several melodic lines. fMRI work on normal listeners implicates the superior temporal gyrus and inferior frontal gyrus in the recognition of a target melody interleaved with distracting tones. Neurological studies of amusia after cortical damage describe associated deficits in pitch perception but there is little information about deficits in the segregation of elements of music. One report described a patient with difficulty perceiving “the whole” of a piece in music following a right-hemisphere haemorrhage (Mazzoni et al., 1993), although the patient reported no difficulty distinguishing the different instruments within a piece. No studies have looked systematically at auditory segregation after acquired cortical damage. Patients with congenital amusia, for which a cortical basis is likely (e.g. Hyde et al., 2006), have pitch discrimination deficits, but do not differ from normal controls in classical tests of auditory stream segregation (Foxton et al., 2004). Deficits of generic segregation and grouping processes relevant to auditory scene analysis as well as deficits of auditory spatial processing have been described in patients with Alzheimer’s disease (Goll et al., 2012; Golden et al., 2015a, 2015b), sparing early auditory cortex (Kurylo et al., 1993) and with cortical substrates in postero-medial and lateral temporo-parietal cortices (Buckner et al., 2009; Seeley et al., 2009; Zhou et al., 2010; Warren, Fletcher & Golden, 2012). However, a study more specifically addressing music streaming did not detect a deficit in this group (Golden et al., 2017).

Here, we report a case of a young woman who experienced a single embolic infarct affecting high-level auditory cortex, who reported a dramatic change in her ability to understand speech in noisy places, and to follow separate lines of music. This case is rare because the patient was only 33 years old and we have no evidence to suggest that she had peripheral damage that could contribute to higher-level processing impairments, or other processes such as small vessel disease affecting the brain as commonly occurs in older subjects. We were able to carry out detailed psychophysics to describe the nature of her auditory processing deficits following the stroke.

## 2. Case Report

The patient was a healthy 33-year-old woman with a history of misophonia, but no history of hearing difficulties other than recurrent ear infections as a child. She was educated to post-graduate level. She was musical and had learnt to play the piano between the ages of 7 and 11 years.

The patient experienced hearing symptoms coincident with a right hemisphere stroke manifest as sudden-onset weakness and loss of sensation in the left arm and leg associated with nausea, vomiting and collapse. This was felt to be due to a paradoxical embolus associated with a deep vein thrombosis that passed from the right to the left heart through an atrial defect. She reported becoming aware of hearing difficulties on the day of her event.

She reported difficulty hearing music through her headphones: the volume was adjusted to the highest setting, but she could only hear part of the music. However, this difficulty was only transitory. Shortly afterwards, she attended a family gathering and quickly realised she was struggling to identify who was speaking. She also reported difficulties processing speech in group situations when there was background noise—which became worse in environments with prominent echo. She also reported finding it difficult to identify emotion in other people’s voices and to identify when someone was asking a question based on inflexion.

She reported no difficulty recognising musical tunes, but commented that music sounded different after her stroke. Familiar music sounded slow and was frustrating to follow. The last part of a song or lyric appeared to merge into the next. She found it easier to listen to music that was played with a single instrument or only vocals. When vocals and background music were present, she could identify the vocals but was unable to hear the background music.

She had previously enjoyed A Capella music (she had friends in a group), but now found it difficult to pick out the different voices. She also struggled to identify emotion in music. She reported that, in general, familiar sounds (such as a running tap) sounded distorted, and described them as sounding ‘tinny’ and ‘echoey’. She described difficulty localising sounds— particularly traffic sounds when crossing the road. She also reported that she had difficulty ‘tuning into’ sounds, such as the sound of her alarm clock.

The patient had a history of misophonia (Kumar et al., 2014; Schröder, Vulink, & Denys, 2013), which began during her childhood (age 12) and was mainly triggered by the sounds of her father eating and sniffing. In adulthood, she described similar misophonic reactions to sounds made by her husband, such as ‘clicking’ sounds when he spoke. After her stroke, she perceived more ‘clicking’ sounds in speech. Breathing noises—which had not bothered her previously—also triggered misophonic reactions. In general, she found that a wider variety of sounds triggered misophonic reactions (for example, the distorted sound of the running tap), and she experienced misophonic reactions more intensely. Other triggers included the sound music from headphones worn by others and the sounds made by people typing at work. She had numerous misophonic episodes, to the point where she described it as ‘unpleasant to exist’.

## 3. Methods

We assessed the patient on four visits to the Royal National Throat, Nose and Ear Hospital (University College London Hospitals NHS Foundation Trust), which took place 9, 10, 14, and 22 months after the stroke. She reported that her symptoms were relatively stable during this period of time—which included problems listening to speech when other sounds were present, a lack of enjoyment for music, and intense misophonia.

The patient underwent a standard protocol of audiological and cognitive assessments and MR testing. We also administered extended auditory psychophysics tests, based on her reported symptoms.

The patient provided written consent for publication of this case report.

### 3.1. MR testing

The patient had a whole brain MRI performed on a 3T Siemens Skyra scanner 3 months after the stroke. The acquisition techniques included T1-weighted 3 dimensional spin echo isometric sequence. The scan acquisition parameters were as follows: repetition time (TR) = 700ms; echo time (TE) = 11ms; number of averages = 2; number of phase encoding steps = 282; acquisition matrix 256 × 256; flip angle = 120; contrast agent = 12 ml gadolinium (Dotarem®).

### 3.2. Audiological testing

A routine audiological test battery was performed, including tympanometry (performed using a GSI 33 Middle Ear Analyser, Grason Stadler) and pure tone audiometry (performed using a GSI 61; Grason Stadler). Normal tympanometry—recorded at 226 Hz—was determined by a sharp single peak, middle ear pressure between –50 and +50 daPa, and compliance of 0.3–1.6 (British Society of Audiology, 2013). Pure tone thresholds were measured at .25, .5, 1, 2, 3, 4, 6, and 8 kHz in each ear. Thresholds ≤ 20 dB HL were considered to be within the normal range (British Society of Audiology, 2004).

We measured transient evoked otoacoustic emissions (TEOAEs) in both ears using the ILO88/92 Otodynamic Analyser with a standard setup (Kemp, Ryan, & Bray, 1990). The presence of normal TEOAEs at 500–4000 Hz was determined by overall signal-to-noise ratios ≥ 6 dB and waveform reproducibility of > 70% (Hurley & Musiek, 1994). We also measured contralateral suppression for TEOAEs using a broadband masker. To calculate suppression, we subtracted the TEOAE amplitude when it was measured in the presence of the contralateral masker from the amplitude measured without contralateral stimulation. Suppression ≥ 1 dB is considered to be within the normal range (Coelho, Ceranić, Prasher, Miller, & Luxon, 2007).

The Speech, Spatial and Qualities of Hearing Scale (SSQ) (version 3.1.2; Gatehouse and Noble, 2004) was used to assess the patient’s perceived auditory disability. The questionnaire contains 14 Speech items, 17 Spatial items, and 19 Qualities items. Each item uses a 10-point rating scale, where higher ratings indicate better self-reported abilities. Speech scores < 6.84, Spatial Scores < 6.14, or Quality scores < 8.18 indicate perceived disability (Demeester et al., 2012).

### 3.3. Additional tests

#### 3.3.1. Gaps in noise

The gaps in noise test (Musiek et al., 2005) was used to assess within-channel temporal resolution in each ear. On each trial, flat-envelope broadband noise was presented for a duration of 6 seconds. The noise contained 0–3 silent gaps, which each had a duration of 2–20 ms. The patient was instructed to press a button as quickly as she could whenever she perceived a gap. Each test contained 60 gaps in total (6 gaps per duration; durations of 2, 3, 4, 5, 6, 8, 10, 12, 15, and 20 ms), and we used different stimulus sets for each ear.

The gaps in noise test was played from a compact disk on a Sony CD Player, which was presented monaurally through a GSI 61 diagnostic audiometer to TDH-39 matched earphones. The test was conducted in a quiet room and sounds were presented at 60 dB SPL.

The gap detection threshold was calculated as the shortest gap duration at which the patient was able to correctly perceive the gap at least 4 out of the 6 times it was presented. The detection threshold is considered to be within the normal range if it is 6 ms or shorter (Musiek et al., 2005).

#### 3.3.2. Pitch discrimination

Pitch difference limens at 1000 Hz were based on the procedure reported by Foxton et al. (2004). Figure 1A shows a schematic of the trial structure. On each trial, participants were presented with two pure tone pairs. Each pure tone lasted 200 ms and was gated by a 10 ms raised-cosine ramp. Within each interval, there was a silent gap of 100 ms between tones, and the two intervals were separated by 400 ms. The two pure tones in one of the two pairs were both 1000 Hz (50% interval 1, 50% interval 2). For the other pair, one tone was higher and one tone was lower than 1000 Hz. On each trial, the patient was asked to identify whether the first or the second interval contained two tones that differed in frequency. The frequency difference began at ±20% and we used a 1-up 2-down procedure (Levitt, 1971) to estimate the 70.7% threshold. The step size ratio was √2 and the inter-trial interval was 0.8–1.2 seconds. The procedure stopped after 6 reversals and we repeated the full procedure twice.

**Figure 1.**
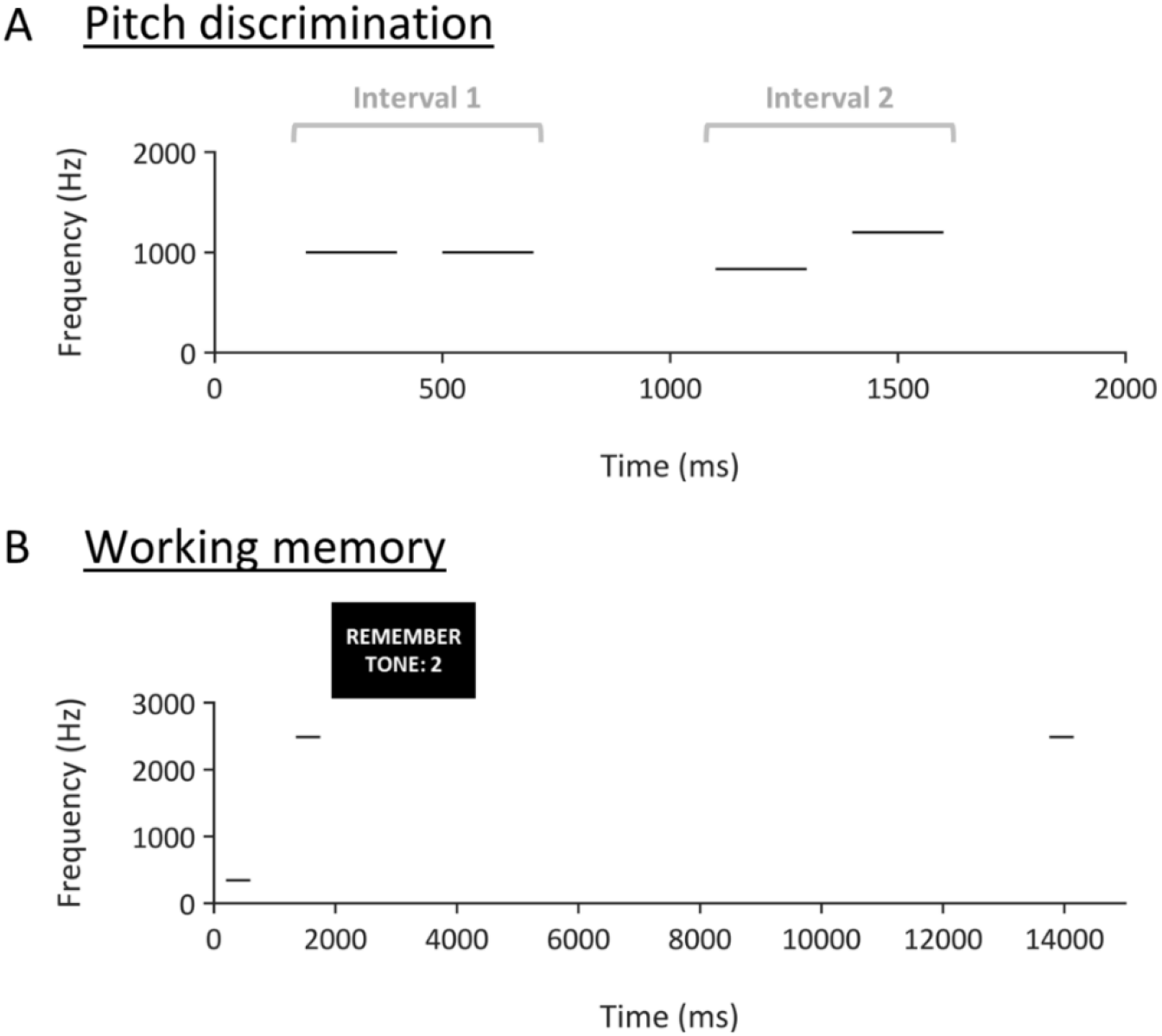
A. Schematic of pitch discrimination task. Each line represents a pure tone with a duration of 200 ms. B. Schematic of working memory task.

**Figure 2.**
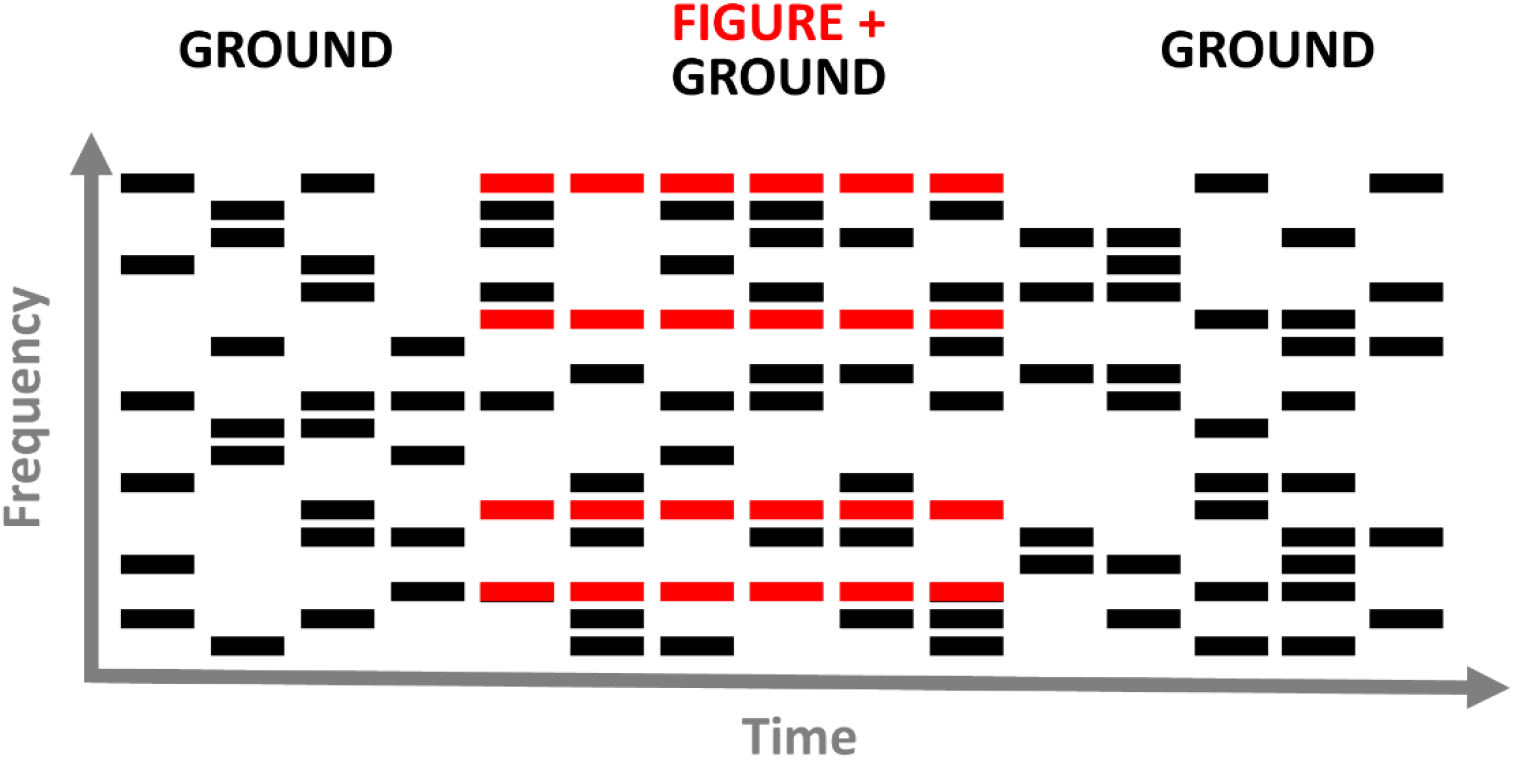
Schematic of a stochastic figure-ground stimulus with a duration of 6 chords and a coherence of 4. Each bar represents a tone of 50 ms duration. Red bars belong to the ‘figure’ and black bars belong to the ‘ground’. Note that an excerpt of 15 chords are displayed here, whereas the entire stimulus lasted for 40 chords.

The pitch discrimination task was presented using MATLAB (R2017a). Stimuli were presented diotically through circumaural headphones (Sennheiser HD 380 Pro) connected to an external sound card (ESI Maya 22 USB) and were presented at 75 dB A.

We calculated pitch difference limens as the median of the last 4 reversals in each procedure. Cut-off values were calculated as 2 standard deviations from the mean from control data in Foxton et al. (2004): 0.36 semitones.

#### 3.3.3. Frequency pattern

The frequency pattern test measures the ability to discriminate three-tone sequences containing mixtures of high (1122 Hz) and low (880 Hz) frequency tones. Each tone lasted 150 ms and the inter-tone interval was 300 ms. After each sequence, the patient was instructed to repeat the pattern she heard (e.g., high-low-high). We presented 30 trials to each ear, including 3 practice sequences.

The frequency pattern test was played from a compact disk on a Sony CD Player, which was presented monaurally through a GSI 61 diagnostic audiometer to TDH-39 matched earphones. The test was conducted in a quiet room and tones were presented at 60 dB SPL.

Performance on the frequency pattern test was calculated as the percentage of patterns reported correctly in each ear. Scores ≥ 78% are considered to be within the normal range (Musiek, 1994).

#### 3.3.4. Auditory working memory

We measured working memory for pitch using the trial structure illustrated in Figure 1B. On each trial, a high tone (312, 342, or 534 Hz) and a low tone (2488, 2643, or 2790 Hz) were presented (in either order) with an inter-stimulus interval of 750 ms. Each tone lasted 400 ms and was gated by a 10 ms raised-cosine ramp. After the two tones had ended, the patient saw a visual cue that instructed her to remember the pitch of the first or the second tone. After a 12 second delay, a single pure tone was presented that was either identical (50% of trials) or different to the cued tone. When the tone was different, it was 10% higher or lower in frequency than the original. The patient was asked whether the tone was the same or different to the cue, and responded by pressing a button on a computer keyboard. We presented 24 trials with an inter-trial interval of 3.25–4.25 seconds. Prior to the main task, the patient completed 8 practice trials with feedback, which were not included in the analysis.

The auditory working memory task was presented using MATLAB (R2017a). Stimuli were presented diotically through circumaural headphones (Sennheiser HD 380 Pro) connected to an external sound card (ESI Maya 22 USB) and were presented at 75 dB A.

We calculated the percentage of trials with correct responses. Normative data collected from 22 subjects (Dheerendra, Kumar, and Griffiths, unpublished) showed a mean performance of 79.0% (standard deviation = 8.8). Therefore, we define normal performance as performance ≥ 61.4% (i.e., within 2 standard deviations of the mean).

#### 3.3.5. Auditory figure-ground

We measured basic auditory segregation with stochastic figure-ground stimuli, similar to those used by Teki et al. (2013). Stimuli consisted of 40 50-ms chords with 0 ms inter-chord interval. Each chord contained multiple pure tones that were gated by a 10-ms raised-cosine ramp. The background comprised 5–15 pure tones at each time window; the frequencies were selected randomly from a logarithmic scale between 179 and 7246 Hz (1/24th octave separation). The background lasted 40 chords (2000 ms). The figure started on chord 15–20 of the stimulus. We used figure coherence levels of 4, 6, 8, and 12 components and durations of 4, 6, and 8 chords (i.e., 200, 300, and 400 ms). The frequencies of the 4–12 figure components were selected randomly, but with an additional requirement that the figure frequencies were separated by more than one equivalent rectangular bandwidth (ERB). The frequencies of the figure were the same at adjacent chords. For half of stimuli, there was no figure in the stimulus; to ensure that figure-present and figure-absent stimuli had the same number of elements (and therefore the same amplitude), figure-absent stimuli contained an additional 4, 6, or 8 components of random frequencies, which had the same onset and duration as the figures in figure-present stimuli. The patient’s task was to decide whether a figure was present or absent on each trial. We presented 60 trials (5 of each combination of coherence and duration conditions) with an inter-trial interval of .8–1.2 seconds.

The auditory figure-ground task was presented using MATLAB (R2017a). Stimuli were presented diotically through circumaural headphones (Sennheiser HD 380 Pro) connected to an external sound card (ESI Maya 22 USB) and were presented at 75 dB A.

Before the test began, the patient heard 4 examples of figure-absent trials, followed by 2 examples of the figure played alone (which never appeared in the test trials). She then heard 8 examples of figure-present trials. On the first occasion that the patient completed the test, the figure-ground stimuli were presented diotically. The patient also performed the task when the stimuli were presented to the left or right ears alone.

To determine behavioural performance, we calculated d′ (Green & Swets, 1966) with loglinear correction (Hautus, 1995). We compared performance at each coherence and duration level to results reported by Teki et al. (2013: Experiment 1). Given that Teki et al. (2013) did not measure performance at durations as long as 8 chords, we used their duration=7 condition to estimate cut-off values for this condition, which should be a (conservative) lower bound on the true value for duration=8.

#### 3.3.6. Speech-in-noise

##### The patient completed four speech-in-noise tests

The Listening in Spatialized Noise – Sentences (LiSN-S) test assesses the ability to understand spoken sentences in the presence of competing speech that originates from different directions. Stimuli were presented through headphones with a three-dimensional virtual reality auditory environment created by synthesizing the stimuli with head-related transfer functions (Cameron and Dillon, 2007). The patient was instructed to repeat each target sentence, which was presented simultaneously with two distractor stories. The distractor stories could either be positioned at the same location as the target sentence— directly in front of the listener (0° azimuth)—or symmetrically spaced to the left and right of the target (± 90° azimuth). In addition, the distractor voices were either the same as the target voice or different. For each of the four conditions (2 spatial × 2 talker conditions), the level of the target stimulus was adapted in a 1-up 1-down procedure to estimate the signal-to-noise ratio (SNR) for reporting 50% of words correctly. Five different scores are calculated: a low-cue speech reception threshold (SRT), corresponding to the same-location same-talker condition; a high cue SRT, corresponding to the different-location different-talker condition; a ‘talker advantage’ score, corresponding to the difference in dB between thresholds in the same-location same-voice condition and the same-location different-voice condition; a ‘spatial advantage’ score, corresponding to the difference in dB between thresholds in the same-location same-voice condition and the different-location same-voice condition; and a ‘total advantage’ score, corresponding to the difference in dB between thresholds in the same-location same-voice condition and the different-location different-voice condition. Outcome measures are z-scores for the above that are generated by the test programme according to age-specific normative data (Cameron et al., 2011). Normative cut-off values for a 33-year-old are listed in Table 2.

**Table 2.** Performance on each of the tests, displayed next to the normative cut-off values. The final column contains a tick if the patient is within normal limits, and a cross if the patient is outside of normal limits.

| Test | Ear | Test score | Normative cut-off | Within normal range? |
| --- | --- | --- | --- | --- |
| **Gaps in noise** |  |  |  |  |
|  | Left | 10 ms | > 6 ms | ✗ |
|  | Right | 6 ms | > 6 ms | ✓ |
| **Pitch discrimination** |  |  |  |  |
|  | Diotic | First run: 4.42% (0.75 semitones, 75 cents)<br>Second run: 4.27% (0.72 semitones, 72 cents) | > 0.36 semitones | ✗ |
| **Frequency pattern** |  |  |  |  |
|  | Left | 70% | < 78% | ✗ |
|  | Right | 90% | < 78% | ✓ |
| **Auditory working memory** |  |  |  |  |
|  | Diotic | 66.67% | < 61.4% | ✓ |
| **LiSN-S** |  |  |  |  |
| Low cue SRT | Diotic | -0.4 | > 0.5 | ✓ |
| High cue SRT | Diotic | -16.4 | > -10.8 | ✓ |
| Talker advantage | Diotic | 9.5 | < 4.7 | ✓ |
| Spatial advantage | Diotic | 13.3 | < 8.7 | ✓ |
| Total advantage | Diotic | 16.0 | < 9.4 | ✓ |
| **Dichotic digits** |  |  |  |  |
|  | Left | 92.5% | < 95% | ✗ |
|  | Right | 97.2% | < 95% | ✓ |
| **Words in babble** |  |  |  |  |
|  | Left | First test: 7.25 dB<br>Second test: 4.25 dB | > 3.5 dB | ✗ |
|  | Right | First test: 4.00 dB<br>Second test: 1.50 dB | > 3.5 dB | ✓ |
| **Sentences in babble** |  |  |  |  |
|  | Diotic | Run A: -3.25 dB | > 0.1 dB | ✓ |
|  |  | Run B: -2.75 dB |  |  |
| **MBEA** |  |  |  |  |
| Scale | Diotic | 18 | < 22 | ✖ |
| Contour | Diotic | 21 | < 22 | ✖ |
| Interval | Diotic | 16 | < 21 | ✖ |
| Rhythm | Diotic | 28 | < 23 | ✓ |
| Meter | Diotic | 25 | < 20 | ✓ |
| **Golden et al (2017) music battery** |  |  |  |  |
| Pitch (local) | Diotic | .32 | < .73 | ✖ |
| Pitch (global) | Diotic | .69 | <.68 | ✓ |
| Pitch (global-direction-only) | Diotic | .45 | <.48 | ✖ |
| Temporal (local) | Diotic | .64 | <.78 | ✖ |
| Temporal (global) | Diotic | .55 | <.50 | ✓ |
| Timbre | Diotic | 1.00 | < .97 | ✓ |
| Tune streaming | Diotic | .65 | < .74 | ✖ |
| Tune recognition | Diotic | 19 | < 18.7 | ✓ |

We also used the dichotic digits test (Musiek, 1983). On each trial, the patient heard two spoken digits in each ear. Within each ear, the two digits were presented sequentially, and the onsets of the digits were aligned between the ears. The patient was asked to repeat the digits that were presented to both ears. The dichotic digits test was played from a compact disk on a Sony CD Player, which was presented monaurally through a GSI 61 diagnostic audiometer to TDH-39 matched earphones Stimuli were presented at 60 dB SPL. The first three trials were used as a practice and were not included in the scoring. 20 test trials were scored: we calculated the percentage of the digits presented to each ear that were reported correctly. Scores ≥ 95% are considered to be normal.

The words-in-babble test (Spyridakou, Rosen, Dritsakis, & Bamiou, 2019) measured word report separately in each ear. Stimuli were presented monaurally and the patient was asked to repeat each word. In total, 25 monosyllabic words were presented simultaneously with 20-talker babble. An adaptive procedure was used to determine the signal-to-noise ratio (SNR) corresponding to the 50% threshold. The masker level was fixed (65 dB SPL), and the intensity of target was adapted according to the SNR. The SNR began at 12 dB, and was adapted in 6 dB increments, which decreased to 2 dB in 1 dB increments at each reversal. The test was conducted in a sound-proof room, and stimuli were presented monaurally through Sennheiser HD 25 headphones, controlled by custom-written Matlab software. The test stopped after 25 words, and the threshold was calculated as the mean of the final 6–8 reversals. The procedure was repeated twice in each ear with different word lists. Scores ≤ 3.5 dB are considered to be within the normal range.

We also ran a diotic sentences-in-babble task (Holmes & Griffiths, 2019). Target sentences were from the English version of the Oldenburg International Matrix corpus (HörTech, 2014) spoken by a male native-English speaker. This was a closed-set test, in which the task was to select the 5 words that were spoken on each trial from a list of options (10 options for each word; see Table 1) in any order. The background was 16-talker babble, which began 500 ms before the onset of the target sentence. The babble was extracted from a continuous track lasting 20 seconds; a different segment of the babble was selected on each trial. We adapted the SNR in a 1-up 1-down procedure to estimate the 50% threshold. The procedure began at 0 dB SNR; the step size began at 2 dB and decreased to 0.5 dB after 3 reversals. We used two interleaved runs, and each run terminated after 10 reversals. The sentences-in-babble task was presented using MATLAB (R2017a). Stimuli were presented diotically through circumaural headphones (Sennheiser HD 380 Pro) connected to an external sound card (ESI Maya 22 USB) and were presented at 75 dB A. We calculated the threshold for each run as the median of last 6 reversals. Holmes & Griffiths (2019) found that 97 healthy participants with normal audiometric thresholds scored a mean of −3.1 dB SNR on this task (standard deviation = 1.6). Therefore, typical performance—defined as thresholds within 2 standard deviations of the mean—is ≤ 0.1 dB SNR.

**Table 1.** Words from the English version of the Oldenburg International Matrix corpus. Target sentences in the speech-in-babble task contained one word from each column, which were recorded and presented as full sentences.

| Name | Verb | Number | Adjective | Noun |
| --- | --- | --- | --- | --- |
| Alan | got | three | large | desks |
| Doris | sees | nine | small | chairs |
| Kathy | brought | seven | old | tables |
| Lucy | gives | eight | dark | toys |
| Nina | sold | four | heavy | spoons |
| Peter | prefers | nineteen | green | windows |
| Rachel | has | two | cheap | sofas |
| Steven | kept | fifteen | pretty | rings |
| Thomas | ordered | twelve | red | flowers |
| William | wants | sixty | white | houses |

#### 3.3.7. Music

##### The patient reported a decreased enjoyment of music, so we used two test batteries to assess her musical ability

The Montreal Battery for the Evaluation of Amusia (MBEA) (Peretz, Champod, & Hyde, 2003) was designed as a measure of musical ability for non-musicians. It is not designed for musicians but was used here to screen for striking musical deficits in a musically trained subject. We used five sub-tests. Three of the tests are classified as melodic organization tests: a target and a comparison melody are played sequentially, which are identical except that one of the tones in the comparison melody differs in pitch from the target melody. In the scale test, the pitch does not belong to the same musical key as the rest of the melody, but is consistent with the original melodic contour. In the contour test, the pitch belongs to the correct musical key, but has a different contour direction than the original. In the interval test, the pitch is in the correct contour and musical key, but is a different pitch and therefore could be detected by an interval change relative to the previous tone. For the melodic organization tests, the patient’s task was to decide whether the target and comparison sequences were the same or different on each trial. The position of the different-pitch tone within the melody differed across trials. The final two tests are classified as temporal organization tests. For the rhythm test, a target and comparison melody are played sequentially, and the relative durations of two adjacent tones are different in the comparison melody; although, the meter and number of tones is the same as in the target melody. In the meter test, each trial contains a single melody, and the task is to categorize the melody as either a waltz (triple meter) or a march (duple meter). The tests were conducted in a quiet room and were presented through the speakers of a Dell latitude 3450 laptop running MATLAB (R2014A). The patient completed 2 practice trials for each test, followed by 30 test trials. We recorded the number of trials the patient responded correctly. The cut-off scores, defined as performance below 2 standard deviations from the mean of 160 normal subjects, are as follows: 22 for the scale and contour tests, 21 for the interval test, 23 for the rhythm test, and 20 for the meter test (Peretz et al., 2003).

The second music battery was developed by Golden et al. (2017). It aims to test musical perception while minimizing working memory load. The battery contains 5 tests. Three of the tests required the patient to detect deviant tones within a sequence; she was asked to press a button as soon as the deviant tone occurred. In the timbre deviant task, melodies were musical scales, and deviant tones had a different spectral envelope than the other tones in the melody. In the pitch deviant task, melodies were arpeggios or Alberti bass sequences; deviants were either classified as local (they fit the contour but had the wrong pitch), global (they were in the opposite direction to the melodic contour and did not belong to the set of pitches contained within the pattern) or global-direction-only (they matched the pitch of one of the other tones in the repeated pattern, but were in the opposite direction to the melodic contour). Responses that occurred ≤ 1.5 seconds after the onset of the deviant tone were classified as correct. In the temporal deviant task, all of the tones had the same pitch, and deviants were either local violations (e.g., two tones to replace a single longer-duration tone) or global violations (e.g., an extra beat in the bar, representing a deviation from the time signature). Responses that occurred ≤ 2.0 seconds after the onset of the deviant tone were classified as correct. In the tune streaming test, the patient heard 20 melodies that were either highly familiar or novel (10 of each type; novel melodies were pseudo-reversed versions of the familiar melodies) against a melodic background containing two lines of music. She was asked to identify whether the embedded melody was familiar or unfamiliar. Finally, as a baseline for the tune streaming test, a tune recognition test was delivered, in which the same 20 familiar and novel melodies were presented alone. We counted the number of trials the patient responded correctly. For both of these two tasks, the patient was asked to decide whether the tune was familiar or unfamiliar. There were no practice trials, but the tests were explained using visual aids, as in Golden et al. (2017). The music battery was presented using MATLAB (R2017a). Stimuli were presented diotically through circumaural headphones (Sennheiser HD 380 Pro) connected to an external sound card (ESI Maya 22 USB) and were presented at a comfortable listening level. For the deviance detection tests, we calculated the corrected detection score, using the method reported in Golden et al. (2017). Normative cut-off values (based on data reported in Golden et al., 2017, from healthy controls with a mean age of 69.7 years and standard deviation of 4.7 years) are listed in Table 2.

## 4. Results

### 4.1. Lesion

Figure 3 shows MRI transverse slices (T2-weighted and fluid-attenuated inversion recovery [FLAIR]) showing the damaged tissue. Figure 4 shows T1-weighted coronal slices through Heschl’s gyrus and the surrounding area. Mature signal and volume changes consistent with infarction were evident in the right cerebral hemisphere, involving the cortical and subcortical regions of the inferior parietal lobule, the parietal operculum, the posterior aspect of the superior temporal gyrus, and part of the postcentral gyrus. The damage affected the temporo-parietal junction into planum temporale and there was some impingement onto Heschl’s gyrus (HG), although medial HG appeared to be spared. This was determined by visual inspection of the anatomical images and by projecting a map of right Te1.0 (Morosan et al., 2001; SPM Anatomy Toolbox version 2.2c: Eickhoff et al., 2005) onto the T1 image, after normalisation using SPM12 (Wellcome Centre for Human Neuroimaging, London, UK). A small area of the posterior insula was also affected. There was no evidence of previous haemorrhage in this region, or any other.

**Figure 3.**
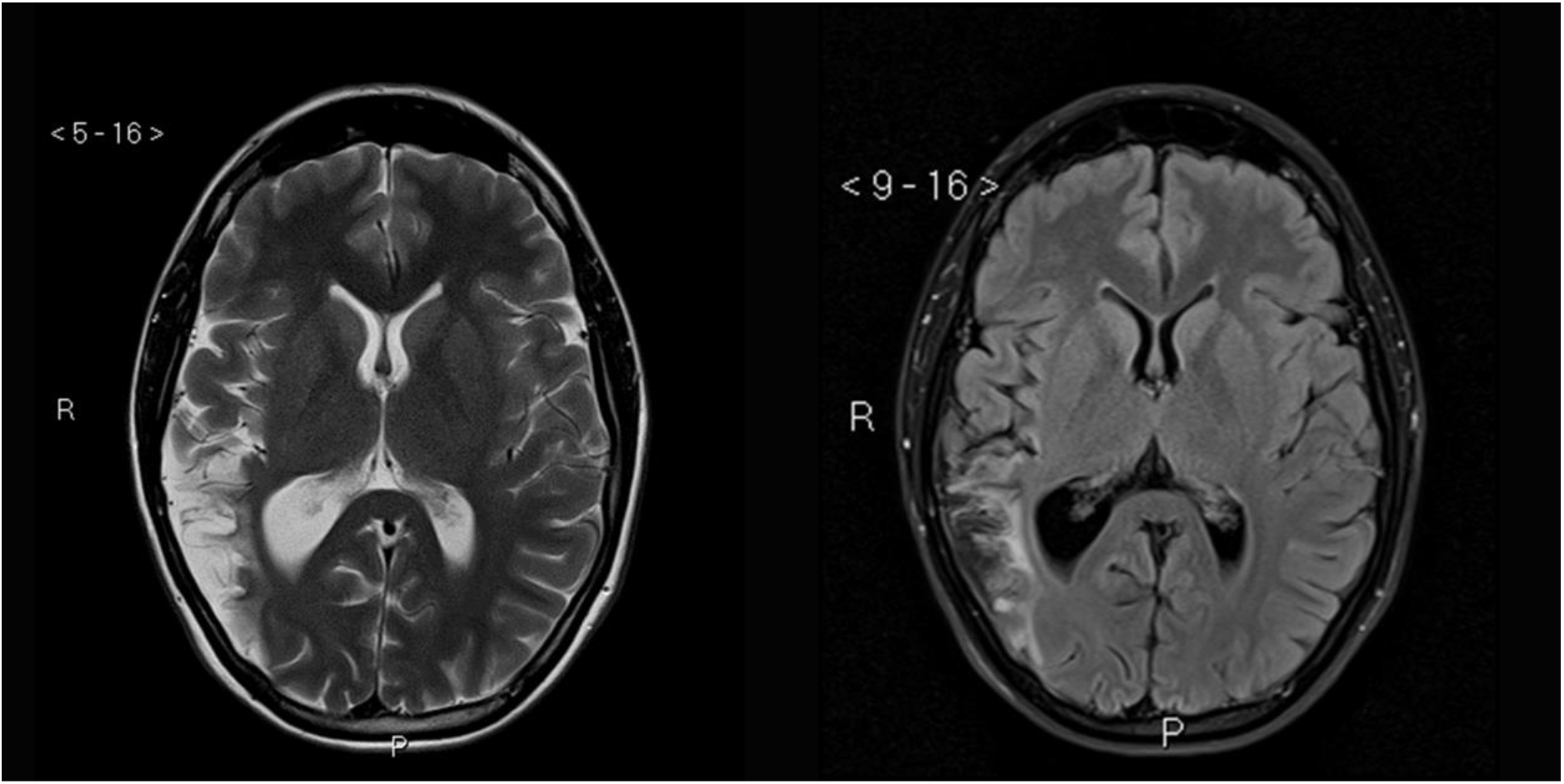
Axial MRI slices demonstrating a single infarct in the territory of the inferior division of the right middle cerebral artery. Left: T2-weighted. Right: Fluid-attenuated inversion recovery (FLAIR). R: Right; P: Posterior.

**Figure 4.**
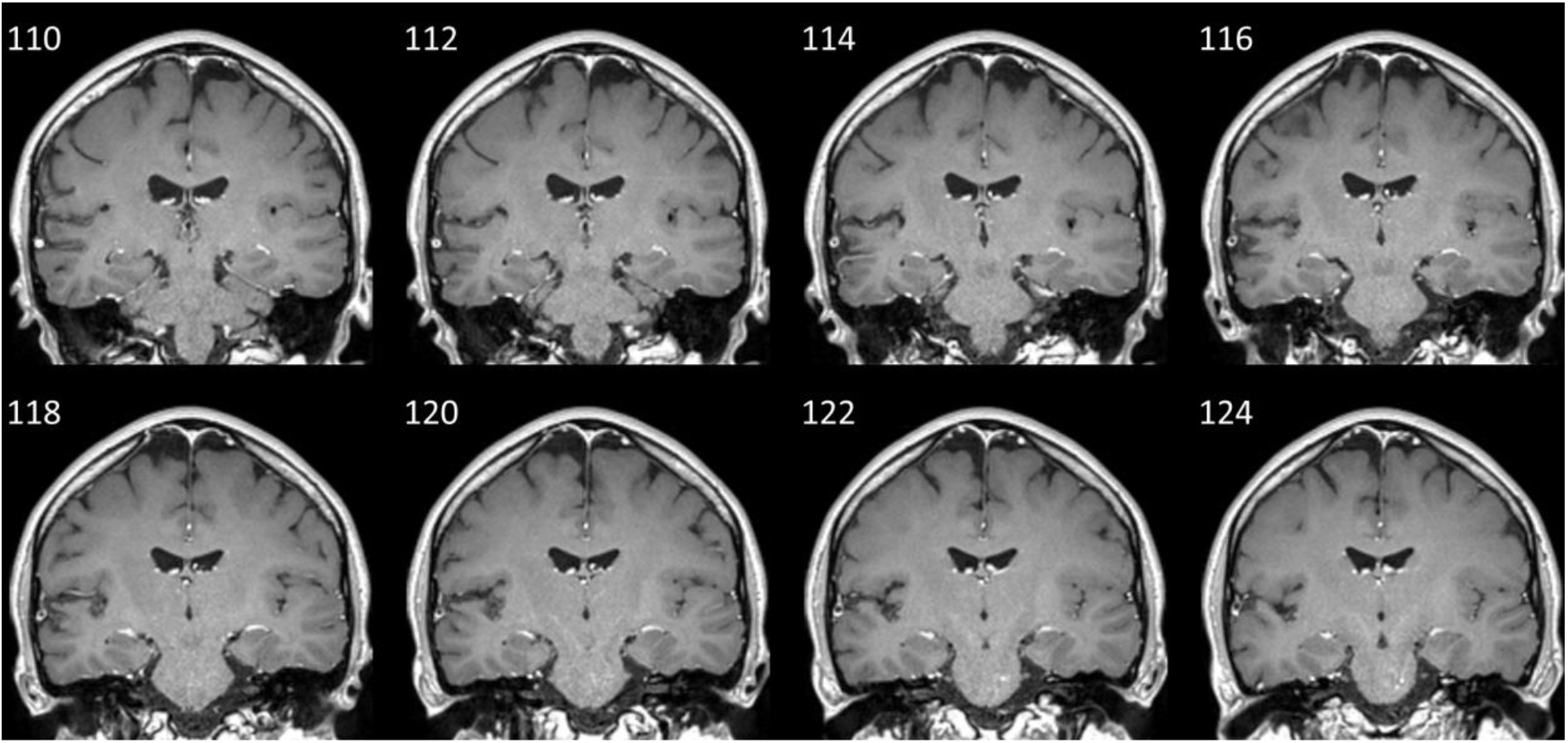
Coronal slices of the T1-weighted MRI image, separated by 2 mm, demonstrating signal changes in the region of auditory cortex lateral to medial Heschl’s Gyrus (HG) in the right superior temporal plane. Displayed in radiological convention, with the right of the brain on the left side of the image. Y co-ordinates (mm) are displayed in the upper left of each slice.

### 4.2. Subject reports

SSQ scores indicated perceived disability in all three domains (Speech = 4.29, Spatial = 2.24, Qualities = 2.32).

### 4.3. Audiological testing

The tympanometry traces for both ears had normal sharp single peaks. The patient had normal middle ear pressure and compliance in both ears.

Figure 5 shows the results of pure-tone audiometry. The patient had normal pure-tone thresholds, which were < 20 dB HL at all of the frequencies we tested.

**Figure 5.**
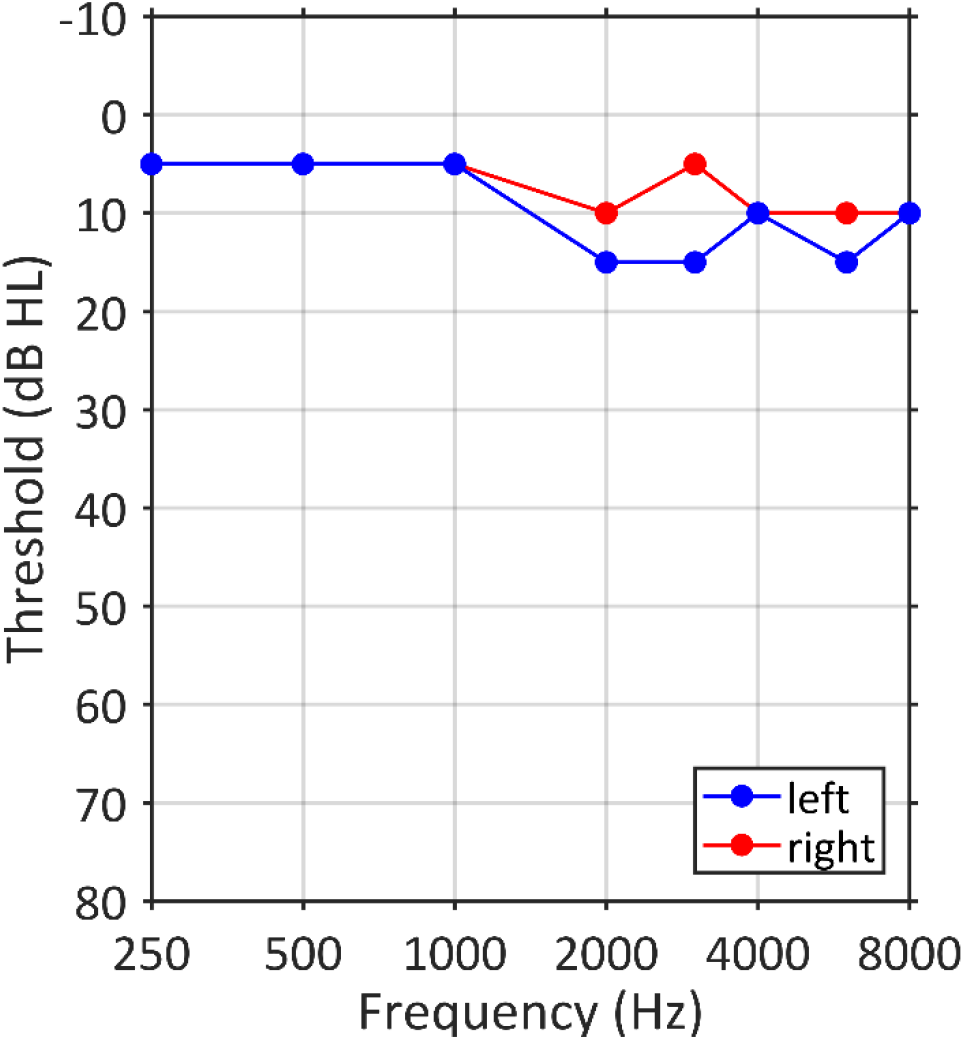
Pure-tone audiogram in the left and right ears.

TEOAE amplitudes were normal (6.4 dB SNR in the left ear, and 9.5 dB SNR in the right ear) and waveform reproducibility was good (83% in the left ear, and 90% in the right ear). Suppression for the left ear (2.5 dB) was within the normal range, whereas it was slightly below in the right ear (0.5 dB).

### 4.4. Additional tests

Tables 2–3 list the patient’s scores for each test, next to the normative cut-offs.

For the left but not right ear, the patient showed atypical performance on the gaps-in-noise and frequency pattern tests. Her scores on the diotic pitch discrimination test were atypical. Her performance on diotic auditory working memory was below average but within normal limits.

For the auditory figure-ground test (Table 3), the patient showed the expected pattern of better performance when the figure had a longer duration and greater coherence (Figure 6). Diotic performance was below average at coherence levels of 4 and 6, but within normal limits. Impairments were present for the left ear at a coherence level of 8 for durations of 6 and 8 chords.

**Table 3.** Sensitivity (d′) for the diotic and monaural conditions of the figure-ground task. Stars indicate scores below the cut-off. Normative cut-offs were estimated from Teki et al. (2013), using a criterion of 2 standard deviations below the mean. There were no normative data for duration = 8 conditions (indicated by the dagger symbol), so the cut-off values are based on the closest condition from Teki et al. (duration7) and therefore can be considered as (conservative) lower bounds on the true value.

| Condition | Diotic | Monaural<br>(left) | Monaural<br>(right) | Normative<br>cut-off |
| --- | --- | --- | --- | --- |
| Coherence = 4 |  |  |  |  |
| Duration = 4 | 0 | 0 | .71 | -.03 |
| Duration = 6 | .71 | 1.17 | 2.06 | .02 |
| Duration = 8 | 1.17 | 1.17 | 2.06 | -.11<sup>†</sup> |
| Coherence = 6 |  |  |  |  |
| Duration = 4 | 0 | 0 | 1.17 | -.05 |
| Duration = 6 | 1.59 | .88 | 2.06 | .50 |
| Duration = 8 | 1.59 | 1.59 | .42 | .16<sup>†</sup> |
| Coherence = 8 |  |  |  |  |
| Duration = 4 | 2.77 | .88 | 2.77 | .09 |
| Duration = 6 | 2.77 | 0* | 2.06 | .30 |
| Duration = 8 | 2.77 | 1.35* | 2.06 | 1.75<sup>†</sup> |

**Figure 6.**
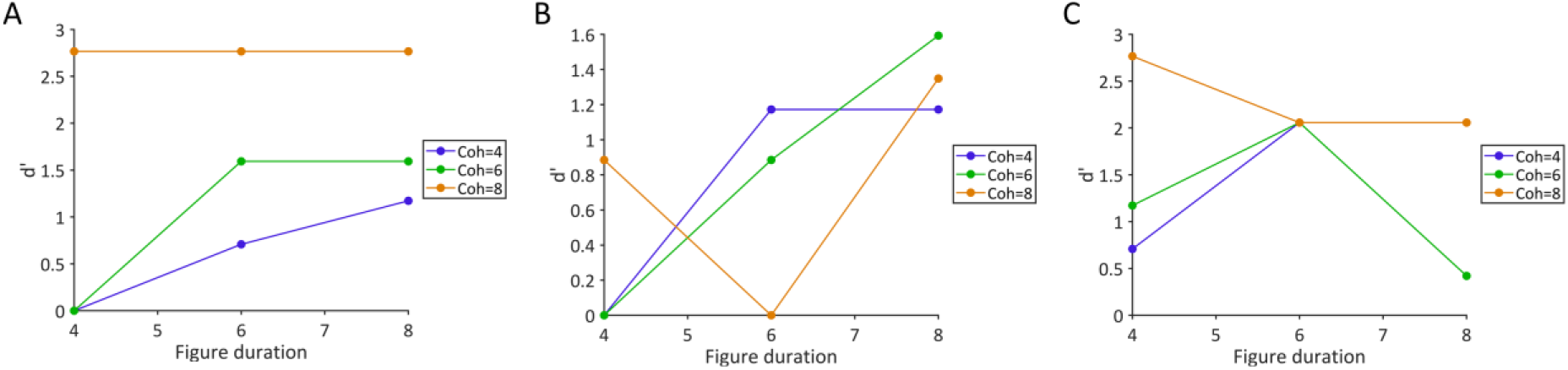
Sensitivity (d′) on the figure-ground task: (A) Diotic presentation; (B) Monaural left ear presentation; (C) Monaural right ear presentation.

Scores on the LiSN-S test were all within the normal range, as were the scores for the diotic speech-in-babble task. For the dichotic digits test, performance was normal in the right ear and below the cut-off in the left ear. For the words-in-babble test, scores for both ears were outside the cut-off in the first presentation, but on the second presentation, the right ear was within normal limits. Thresholds for the left ear were outside the cut-off in both presentations.

The patient showed impairments on the Scale, Contour, and Interval tests of the MBEA. She performed within normal limits on the two temporal organization tests. The patient’s scores fell outside of the normative cut-offs on most of the sub-tests of the Golden et al. (2017) music battery, including those assessing pitch, temporal, and tune streaming. Her score on the global aspect of the pitch test was just inside the normal range, as was performance for the global aspect of the temporal test. Performance on the timbre and tune recognition sub-tests were normal.

## 5. Discussion

In summary, we report the case of a young woman who experienced a domain-general deficit in auditory segregation following a right hemisphere infarction, which affected the inferior parietal lobule, posterior insula, and auditory cortex including planum temporale (PT), but spared medial HG. The deficit was expressed as atypical performance for words-in-noise, music streaming, and figure-ground perception—despite intact peripheral function, working memory, and recognition of familiar melodies. In other words, segregation between objects was impaired when competing sounds were present, despite preserved within-object analysis when object features were tested in isolation. Auditory scene analysis was also somewhat impaired: the patient showed atypical performance on the frequency pattern test, and on musical pitch and temporal deviance detection tasks. Her deficits were most pronounced for sounds presented to the left ear, which is consistent with a right hemisphere lesion (Bamiou et al., 2006). We attribute the impairments in auditory segregation to damage to non-primary auditory cortex including PT, which—in healthy subjects—has been implicated in the types of segregation that were impaired in this patient.

The patient’s descriptions are consistent with immediate auditory deafness, which evolved into auditory agnosia—which is not uncommon (Mendez & Geehan, 1988)—and a worsening of pre-morbid misophonia. The most striking aspect of her agnosia is a deficit in segregation in the speech and musical domains which has not previously been systematically studied.

### 5.1. Word segregation impairment

The patient reported a change in speech perception following her stroke, and reported particular difficulty understanding speech in noisy rooms—when several people were talking at the same time. Interestingly speech-in-noise performance was normal when speech was spatially separated (LiSN test) or presented diotically to both ears (diotic sentences-in-babble thresholds). Whereas, the dichotic digits test and monaurally conducted words-in-babble test both showed deficits for speech presented to the left ear.

This is not a simple case of word deafness, because the patient had no difficulty understanding speech when it was presented diotically or with spatial separation. She was also able to engage in conversation with no difficulty in one-on-one settings. Instead, she specifically found the addition of background noise to be problematic.

Difficulty understanding speech-in-noise is a common complaint among older people (Gatehouse & Noble, 2004), even when clinical tests of peripheral function are unremarkable. The causes of this difficulty in older people are currently unknown, but could be related to aging of the peripheral or central auditory system that is undetected by common clinical measures. This patient is unusual because she was young and we have no reason to suspect peripheral dysfunction. Previous studies have demonstrated that speech-in-noise understanding varies widely among young people with normal hearing (Holmes & Griffiths, 2019), and the neural substrate is likely at early stages of the auditory cortical hierarchy (Holmes et al., 2019). Putative core auditory cortex was spared in this patient, although posterior HG—which Holmes et al. (2019) associated with difficulty in both speech-in-noise and figure-ground perception—was damaged and may, therefore, be related to the patient’s impairments. It is worth noting, however, that the effects reported by Holmes et al. (2019) were strongest in left auditory cortex, whereas this patient’s lesion was confined to the right hemisphere.

### 5.2. Music segregation impairment

Consistent with a generic segregation problem, the tune streaming test of the Golden et al. (2017) music battery was outside of normal limits, despite near-perfect recognition of the same famous tunes presented alone. Both tests required the patient to recognise a target melody; the only difference was that the tune streaming test also contained simultaneous musical tones at different pitches. Intact recognition of famous tunes is not uncommon in cases of right hemisphere lesions (Peretz & Zatorre, 2005), and means this is not an associative form of auditory agnosia. Normal recognition of famous tunes presented alone also rules out several other possible explanations for impaired tune streaming performance: the deficit cannot be because poor working or long-term memory prevented tune recognition, and it cannot be related to impaired pitch or temporal sequencing. Instead, this pattern of results suggests a specific impairment when other musical notes were played simultaneously, mirroring the speech-in-noise segregation problem described above.

It is worth noting that the musical tests we used were designed for non-musicians and the patient had a musical background. In addition, normative values for the Golden et al. (2017) battery were based on data from much older adults and therefore these comparisons likely underestimate the patient’s deficits. Therefore, the fact we found deficits in these tests is particularly striking.

### 5.3. Segregation impairment at a fundamental level

A more abstract task that requires the segregation of pure tone elements—stochastic figure-ground perception—showed a deficit in the left ear. The deficit was most pronounced for the conditions that are usually most salient for healthy subjects: conditions in which the figure contained more frequency elements and had a longer duration. The figure-ground deficit is consistent with the idea that both speech and music segregation problems observed in this patient could arise from impairments in segregation processes that operate at a fundamental auditory level.

Previous descriptions of musical and speech agnosia support the idea that these rarely occur in isolation. More than half of amusic patients also have deficits in speech perception, and approximately one third have difficulties recognising environmental sounds (Stewart et al., 2006). Most previous case studies have chosen to focus on one particular domain, meaning co-occurrence of deficits has probably been underreported (Oppenheimer & Newcombe, 1978). A compelling explanation for common deficits across domains is that these can be caused by deficits in spectrotemporal analysis causing apperceptive auditory agnosia in multiple domains. This argument supports the existence of fundamental deficits in spectrotemporal analysis causing agnosia because of a problem of the analysis of within-object cues. The present report suggests a distinct type of auditory agnosia that is due to the analysis of between-object cues—a segregation deficit—that has not previously been systematically characterised.

The condition that we describe here has some similarities to the visual condition, simultaneous agnosia. In that, patients are unable to segregate complex visual scenes into their component elements. Here, the patient is unable to segregate complex acoustic scenes into their component elements. The visual condition is associated with lesions in the dorsal visual pathway in the parietal lobe and deficits in eye movements and limb movements to visual targets in the periphery: Bálint’s syndrome (Bálint, 1909). The present patient has a lesion in auditory cortex distinct from the lesion in simultaneous visual agnosia. The deficit here is in the segregation of simultaneous objects in time-frequency space as opposed to visual space, and is not accompanied by any symptomatic visual or motor deficits. We suggest the term simultaneous auditory agnosia for the condition, which we argue to be a parallel to simultaneous visual agnosia, in terms of phenomenology and substrate, rather than part of the same syndrome.

In this study, we used tests of fundamental figure-ground analysis to define the deficits in simultaneous auditory agnosia. These figure-ground tests are more abstract and less complex than speech or music, and are devoid of meaning. Yet, they draw on segregation processes that are used by normal individuals to segregate speech from background noise (Holmes & Griffiths, 2019; Holmes et al., 2019). Functional imaging studies of normal subjects based on passive listening or an irrelevant task demonstrate a substrate for these processes that includes auditory cortex (Teki et al., 2016, 2011), and—even though segregation of figure and ground tones occurs during passive listening (Schneider et al., 2018; Teki et al., 2016, 2011)—an effect on the process of attention has been demonstrated in several studies (Molloy, Lavie, & Chait, 2019; O’Sullivan, Shamma, & Lalor, 2015). We suggest that the deficit here is in fundamental auditory segmentation that affects multiple auditory cognitive domains based on the demonstrated lesion in auditory cortex.

### 5.4. Left ear deficits

Across all tests, the patient’s deficits were most pronounced in left ear, consistent with a right hemisphere lesion (Bamiou et al., 2006). This is particularly interesting in the context of the auditory segregation deficits described above, because it suggests that high-level segregation processes are partially dissociable for sounds reaching the two ears, despite the fact that information from the two ears is already combined at a subcortical level. Although processing of auditory objects can of course occur after information from the two ears is integrated, this finding suggests that segregation processes operate at least partially on information from one ear: otherwise ear-specific deficits in auditory segregation could not exist.

Influential models of auditory processing have proposed separate streams for auditory processing, suggesting that auditory object information is processed in a ventral pathway, and spatial (Ahveninen et al., 2006; Bizley & Cohen, 2013; Leavitt, Molholm, Gomez-Ramirez, & Foxe, 2011) or spectral motion (Belin & Zatorre, 2000) information is processed, in parallel, in a dorsal pathway. Our findings indicate that auditory object processing in non-primary auditory cortex contains some information about the ear of origin, possibly reflecting a greater integration between different attributes of sound than would be predicted by these parallel processing models.

### 5.5. Pitch processing

The patient performed below normal limits on the frequency pattern test in the left ear and on the musical pitch tests, and had elevated pitch discrimination thresholds. Part of this deficit could be related to impoverished working memory for pitch, which was within normal limits but below average. However, the Golden et al. (2017) music battery aims to minimise working memory load by asking participants to respond as soon as they detect a deviant sound, so poor working memory is unlikely to fully explain impairments in the Golden et al. (2017) tests.

In previous work, lesions to lateral HG and PT have been associated with impaired discrimination of the direction of a pitch change (Johnsrude, Penhune, & Zatorre, 2000; Liegeois-Chauvel, Peretz, Babaï, Laguitton, & Chauvel, 1998; Terao et al., 2006; Tramo, Shah, & Braida, 2002; Zatorre, 1988), and lateral HG has been proposed as a possible ‘pitch centre’ (Stewart et al., 2006). Therefore, the patient’s damage to these auditory cortical regions is consistent with her impairments to pitch sequencing.

The right hemisphere lesion is likely to be of particular relevance: Milner (1962) found that right lobectomies affect pitch pattern discrimination, whereas left lobectomies do not, and Peretz (1990) showed that patients with right cerebral hemisphere strokes could assess neither global nor local information in melodies. Following a review of studies, both Peretz & Zatorre (2005) and Stewart et al. (2006) conclude that studies consistently associate non-primary auditory cortex in the right-hemisphere with processing pitch relationships. Consistent with these previous studies, our patient had a right hemisphere lesion and impairments to both local and global pitch processing, as well as an impairment on the frequency pattern test. This finding is also broadly consistent with neuroimaging data from healthy subjects who are asked to analyse pitch sequences, which has been associated with bilateral—although somewhat right lateralised—activity (Griffiths, Büchel, Frackowiak, & Patterson, 1998; Patterson, Uppenkamp, Johnsrude, & Griffiths, 2002). In addition, activity in right PT has been associated with the perception of melodies (Griffiths & Warren, 2002).

These pitch deficits are unlikely to fully explain the deficit in auditory segregation described above. First, the patient showed deficits in the tune streaming test but not the tune recognition test, which presents the same melodies alone—and this comparison controls for pitch perception within a stream. Second, the patient’s pitch discrimination thresholds were less than one semitone and therefore, pitch recognition is not sufficiently impaired to affect performance on the speech-in-noise, tune streaming, or figure-ground tests we presented here—in which simultaneous sounds were separated by a larger pitch interval. Patients with congenital amusia have been found to show elevated pitch discrimination thresholds, but show normal performance on auditory streaming tests (Foxton et al., 2004)—demonstrating that elevated pitch discrimination thresholds can contribute to deficits in music perception, but are not always accompanied by higher level segregation problems.

### 5.6. Temporal processing

The patient performed within normal limits on the two temporal organization tests of the MBEA, although was atypical on the local (interval) temporal test of the Golden et al. (2017) music battery. The patient also performed outside of normal limits on the gaps in noise test, which relies on within-channel processes (Walker et al., 2003).

Studies of congenital amusia, in which deficits are typically found in pitch but not rhythmic domains, provide support for distinct substrates for pitch and rhythmic analysis (Foxton et al., 2004) and dissociations are reported in the acquired lesion literature (Stewart et al., 2006). In this report we describe a striking deficit in auditory segregation also associated with pitch domain deficits that largely dissociate from temporal domain deficits. This is consistent with a problem with early segregation of streams and processing of pitch patterns within streams requiring right auditory cortex, as opposed to interval and rhythm analysis dependent on widely distributed areas including the cerbellum and basal ganglia (e.g., Teki, Grube & Griffiths, 2012).

### 5.7. Misophonia

One of the symptoms reported by the patient was a worsening of premorbid misophonia. Initially, damage to the insula may be considered consistent with an increased emotional response to sounds. However, the patient’s lesion was confined to the posterior portion of the insula, and we found no damage in anterior areas that have been associated with misophonia in previous work (Kumar et al., 2017). Therefore, the location of the lesion may not explain the patient's heightened misophonia. Given misophonia was present since childhood, we suspect that changes in misophonic reactions after the patient’s stroke were likely related to generic changes in sound perception, given the deficits described in Sections 5.4–5 (above), or to increased stress and anxiety associated with everyday life, rather than to specific structural damage to the insula. Although, we cannot rule this out as a possible explanation.

### 5.8. Conclusion

Here, we show deficits to higher-level segregation processes associated with a right hemisphere lesion affecting non-primary auditory cortex. The deficits were most pronounced for sounds presented to the left ear, and were domain-general—affecting segregation of speech, music, and more basic abstract stimuli. Importantly, impairments segregating speech and music in the presence of other sounds cannot be explained by changes to the simple perception of target sounds alone. We also found some deficits in analysing pitch and temporal patterns. This relatively rare case of a young stroke patient—who had no detectable peripheral impairment—enhances our understanding of higher-level processes that are necessary for segregating simultaneous sounds.

## Acknowledgements

This work was supported by WT091681MA and DC000242-31 to Timothy D. Griffiths. Jason Warren is supported by grants from the Alzheimer’s Society, Alzheimer’s Research UK and NIHR UCL/UCLH Biomedical Research Centre. The funders had no role in study design, data collection and analysis, decision to publish or preparation of the manuscript.

## Author Contributions

E.H., N.U., D.B., and T.D.G. designed the study, interpreted the data, and wrote the manuscript. E.H. and N.U. collected the data. E.H. analysed the data.

